# Topographic variation in neurotransmitter receptor densities explains differences in intracranial EEG spectra

**DOI:** 10.1101/2024.01.09.574882

**Authors:** U.M. Stoof, K.J. Friston, M. Tisdall, G.K. Cooray, R.E. Rosch

**Author notes:** Classification Biological Sciences | Neuroscience.

## Abstract

Neurotransmitter receptor expression and neuronal population dynamics show regional variability across the human cortex. However, currently there is an explanatory gap regarding how cortical microarchitecture and mesoscopic electrophysiological signals are mechanistically related, limiting our ability to exploit these measures of brain (dys)function for improved treatments of brain disorder; e.g., epilepsy.

To bridge this gap, we leveraged dynamic causal modelling (DCM) and fitted biophysically informed neural mass models to a normative set of intracranial EEG data. Subsequently, using a hierarchical Bayesian modelling approach, we evaluated whether model evidence improved when information about regional neurotransmitter receptor densities is provided. We then tested whether the inferred constraints — furnished by receptor density — generalise across different electrophysiological recording modalities.

The neural mass models explained regionally specific intracranial EEG spectra accurately, when fitted independently. Incorporating prior information on receptor distributions, further improved model evidence, indicating that variability in receptor density explains some variance in cortical population dynamics. The output of this modelling provides a cortical atlas of neurobiologically informed intracortical synaptic connectivity parameters that can be used as empirical priors in future — e.g., patient specific — modelling, as demonstrated in a worked example (a single-subject mismatch negativity study).

In summary, we show that molecular cortical characteristics (i.e., receptor densities) can be incorporated to improve generative, biophysically plausible models of coupled neuronal populations. This work can help to explain regional variations in human electrophysiology, may provide a methodological foundation to integrate multi-modal data, and might serve as a normative resource for future DCM studies of electrophysiology.

**Significance Statement:** Understanding the link between measures of brain function and their underlying molecular and synaptic constraints is essential for developing and validating personalised, pharmacological interventions. But despite increasing availability of detailed normative datasets of human brain structure and function — across modalities and spatial scales — translating between these remains challenging.

Using two large normative datasets — intracranial EEG recordings and autoradiographic receptor density distributions — we demonstrate that generative models of these data can link structure to function. Specifically, we show that regional oscillatory neuronal population activity is shaped by the distribution of neurotransmitter receptors. This modelling furnishes an atlas of normative parameter values, which can provide neurobiologically informed priors for in-silico (e.g., Digital Twin) characterisation of normal and disordered brain functioning.

## 1 Introduction

Translation between spatial neural scales is a central challenge in neuroscience, for understanding neurological disorders and for leveraging molecular and cellular insights in clinical applications. Mechanistic descriptions of pathologies, often formulated at the level of synapses and cells (1–4), differ in spatial scale from measurements of human brain function, such as intracranial and scalp electroencephalography (i/EEG) or functional magnetic resonance imaging (fMRI), which are used in guiding treatments of human brain disorders such as epilepsy (5–7). The difficulty in integrating varying levels of description lies in the structural and functional complexity of the brain, which is characterised by interrelated sub-systems and their non-trivial relationship to emergent regional and whole brain functions, and the fact that meso- and macroscale dynamics shape and constrain the activity of the constituent microscale components (8–10).

To illustrate: many aspects of synaptic mechanisms underlying neuron-to-neuron interaction have been revealed both in healthy brains and in disorders (1, 3, 4, 11–13), but how exactly cellular interactions and entangled neurotransmitter systems influence neuronal population activity represents a challenging problem of emergence (14–17). Conversely, phenomena of human brain (dys)function are described at macroscopic scales and measured with tools such as EEG (18–22), but deconstructing these signals to unravel how the underlying interacting neurotransmitter systems contribute, is an ill-defined inverse problem (23).

Addressing this circular explanatory gap has important practical implications. For example, neuropharmacological interventions, such as anticonvulsants or antidepressants, rely on interpreting clinical phenotypes to target specific mechanisms or components of neurotransmission and neuromodulation. And whilst medications’ affinity for types of neurotransmitter receptors is often well-characterised at microscale (24), their effects on meso- and macroscopic dynamics is less accessible — as they depend on a multitude of factors, including balancing mechanisms of non-targeted interacting neurotransmitter systems (25). Conversely, measuring and describing patients’ brain dysfunction, allows only phenomenological, but not aetiological, diagnoses (26, 27). In consequence, the disconnect between levels of description impedes progress towards individualised patient care (28–31).

Despite the complexity, there is evidence that interrelations between synaptic neurotransmitter systems and whole-brain dynamics are identifiable. For example, there are statistical associations between regional cortical oscillatory rhythms measured using magnetoencephalography (MEG) and neurotransmitter receptor expression measured using positron emission tomography (PET) (32), and between multiple microstructural features of the human cortex and MEG signals (33); additionally, regional neuroreceptor profiles in part explain medication-induced changes in fMRI (25). Similarly, spatially distributed neuroreceptor gene expression, assessed through post-mortem microarray profiles, correlate with fMRI activation patterns during cognitive tasks along gradients of microstructural organisation of the human cortex (34).

This illustrates that statistical relationships between neurotransmitter receptor compositions and cortical rhythms emerge at the level of brain regions. At that spatial scale, neurotransmitter receptor densities might serve as proxies for neuronal functioning (35–37). For example, glutamatergic AMPA and NMDA, and GABAergic GABA_A_ receptor densities vary largely across the cortex (37) and show pronounced alterations in pathological tissue (38). Consequentially, these normative, post-mortem, in-vitro autoradiography data of distinct receptors might inform multi-modal in-silico methods aiming to uncover how receptors contribute to brain dynamics, and positive findings in previous statistical (32) and disease pathway models (39, 40) motivate our investigation.

But measuring neuronal population dynamics accurately with non-invasive methods is challenging (41, 42). Alternatively, intracranial EEG — which is now routinely used in epilepsy presurgical assessments to identify pathological networks (5–7, 43) — captures mesoscale activity with high temporal resolution and signal-to-noise ratio (44). Similar to neuroreceptor densities, iEEG signals are region specific: this was revealed by an atlas of putatively normal intracranial recordings remote from epileptogenic tissue, which showed regionally characteristic iEEG power spectral densities (PSD) in canonical frequency bands (45). Therefore, intracranial EEG offers precise measurements of regional brain activity, explainable by the microarchitecture and functioning of neuronal populations.

To investigate the putative neurobiological mechanisms underlying theses measured brain dynamics, computational models of coupled neuronal populations have been widely applied to address a range of neuroscientific questions (23, 41, 46–49). But, although the parameters of these generative models are inspired by biophysical properties of neuronal population, mapping these to the complex interacting neurotransmitter systems in the human brain has remained challenging.

A possible computational approach to integrate both iEEG and neuroreceptor data and to thereby provide neurobiological grounding, is to incorporate the receptor features as prior information to model fitting of electrophysiological recordings. Bayesian approaches such as dynamic causal modelling (DCM) (50, 51) (50–56) allow for this integration of empirically derived priors (57, 58), which enables us to evaluate different hypotheses about how neuroreceptor composition contributes to regional oscillatory signatures (iEEG).

In summary, we used DCM — as a computationally efficient method — to link recordings of regional brain dynamics and neuroreceptor densities through empirically informed generative models of neuronal population function. Through this approach, we provide evidence that neuronal oscillations in the human brain are shaped by regional variation in neuroreceptor densities and, crucially, generated an atlas of neuronal population parameters that can inform future dynamic causal modelling studies.

## 2 Results

### 2.1 Neuronal population models explain regional oscillatory brain dynamics

To describe the genesis of regional cortical population dynamics, we fitted single neural mass models using dynamic causal modelling for each of the 1770 normative iEEG recordings (Figure 1A). The iEEG atlas included stereotactic EEG (SEEG) and electrocorticographic (ECoG) recordings of putatively normal brain regions in >100 individual epilepsy patients (45). These model inversions (without receptor density priors) constituted the first level of the hierarchical approach; it served to estimate initial parameter posteriors and to evaluate model fits across the range of regional brain dynamics.

**Figure 1.**
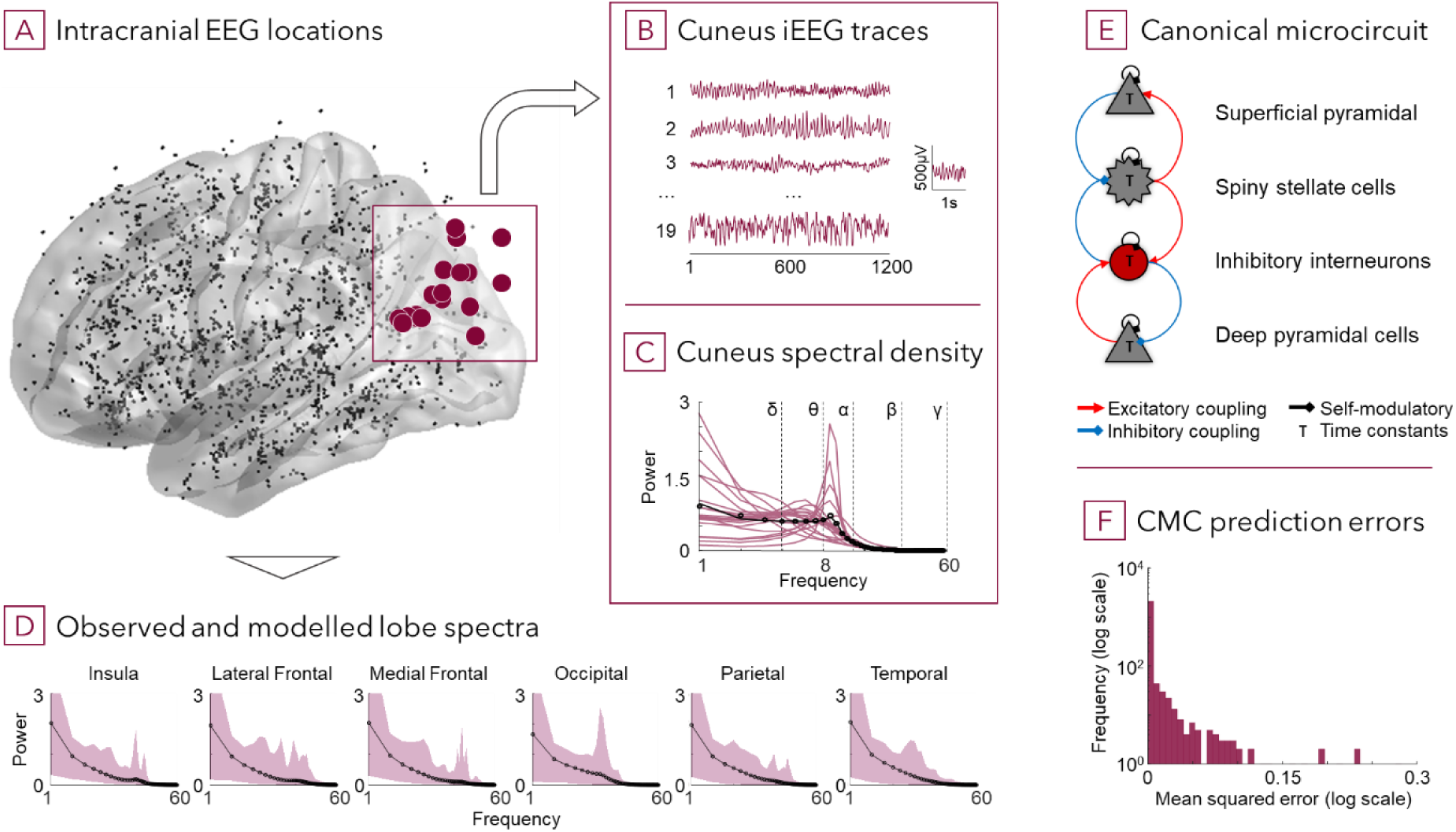
Cortical microcircuit models explain a diversity of regionally specific cortical iEEG patterns. **(A)** iEEG contacts, projected onto a MNI standard space single hemisphere. A total of 1770 traces were included, recorded for a total of 106 patients under awake, eyes-closed conditions without visually apparent pathological features. Red dots highlight cuneus contacts illustrated in subsequent panels. **(B)** Example 5s traces recorded from cuneus contacts. **(C)** Cuneus’ iEEG-derived PSDs for observation (original data), PSDs for individual contacts (grey lines), average of observation over contacts (black line), and average of model (CMC) generated PSDs (black circles). **(D)** PSDs for observation and model averages (black line and circles), and ranges for observation PSDs (red areas) are shown for different cortical lobes and the insula. **(E)** The canonical microcircuit model with four neural populations, which approximate cortical layer-specific activity, was used to model iEEG in the frequency domain. Excitatory (grey) and inhibitory (red) coupling and time-constant posteriors were inverted with Variational Laplace (DCM). **(F)** Most CMC generated PSDs show a good fit and low mean squared error compared to the observation PSDs, only a fraction of models had noticeable deviations, which appear primarily in lower frequencies.

The source-specific, four-population neural mass models (Figure 1E) generated power spectral densities (PSD) (1-60Hz, 1Hz steps) with very good fit for most iEEG recordings with few exceptions (2.4% (42/1770) electrode contacts showed a noticeable error; frequency fit: MSE median = 1.04*10^-6^ and mode = 1.29*10^-11^) (Figure 1F). In addition, the modelled PSDs match previously reported profiles (54–56) and showed characteristic lobe-specific features such as an alpha peak in the occipital lobe (Figure 1C), but also indicated high intra-region and inter-subject variability (Figure 1D).

### 2.2 Variation in neuroreceptor densities explains regional differences in neuronal population dynamics

Next, we tested whether furnishing the regional neural mass models (CMC) with information about neurotransmitter receptor densities improves their fits, as scored with variational (free energy) bounds on log model evidence or marginal likelihood. Since there is no one-to-one mapping between autoradiographic receptor density values and the synaptic parameters of the CMC, this relationship was inferred using a hierarchical parametric empirical Bayes (PEB) approach: this hierarchical model comprised the nonlinear DCM at the first level and a general linear model (GLM) over DCM parameters at the second level. Here, the GLM regressors modelled (between region) variation in receptor densities and the second level inversion estimated the effect of these regressors on first level (within region) parameters. Subsequently, models including different subsets or linear combinations of receptor data were compared to identify the most likely hierarchical model.

First, models that included only receptor density combinations of major excitatory (glutamatergic; AMPA and NMDA) and inhibitory (GABAergic; GABA_A_ and GABA_B_) receptors (Figure 2A) were considered, as these receptor types correspond most directly to the synaptic parameters of the DCM neural mass model. We separately fitted the hierarchical models including regressor combinations of one to four receptor types and compared their evidence using the free energy approximation. The winning model contained AMPA, NMDA and GABA_A_ receptor densities (*ANGa*) with a free energy difference to the next best model of >5, which corresponds to a posterior probability >.99 for *ANGa* over other models.

**Figure 2.**
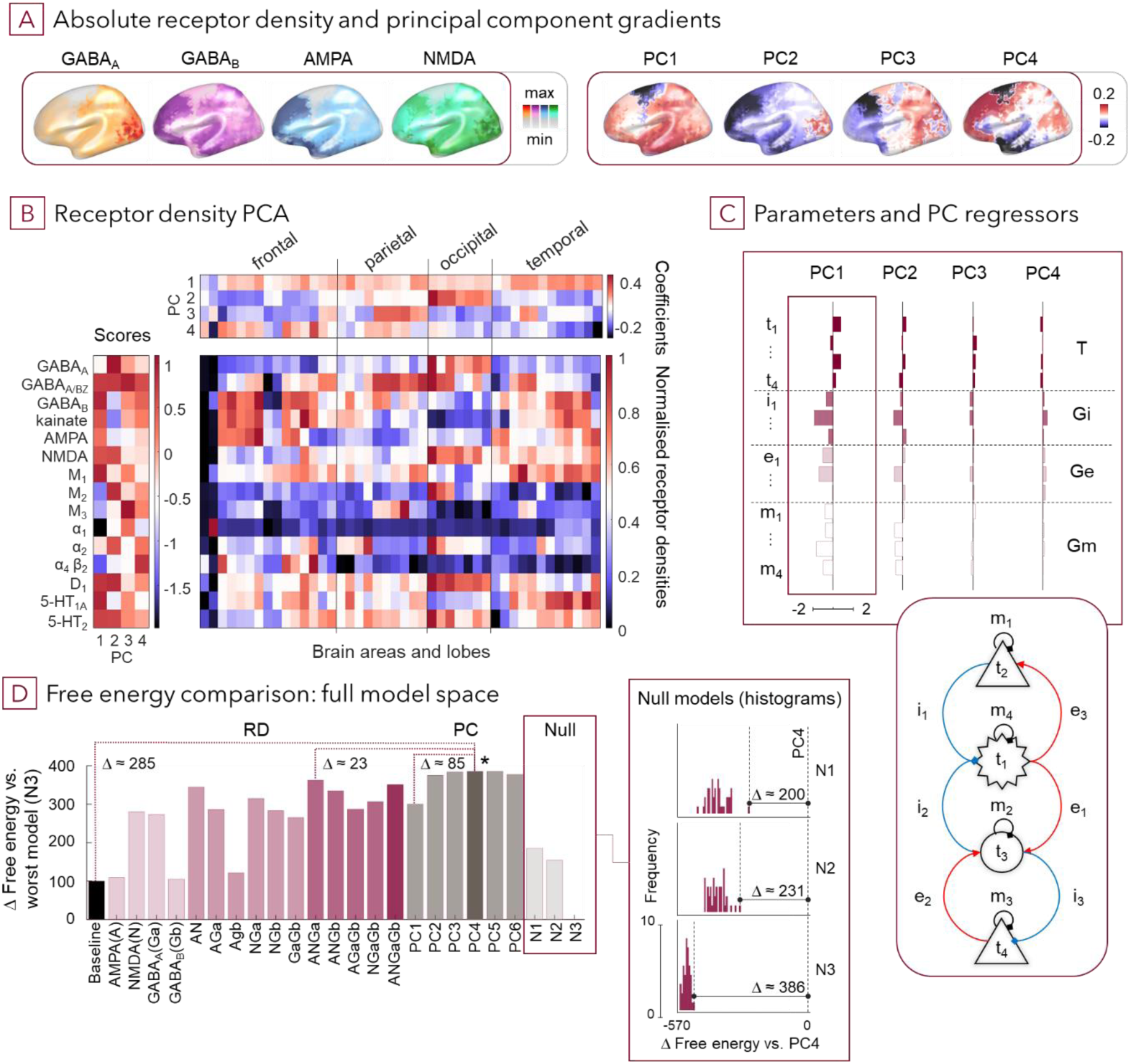
Prior information on the spatial distribution of neurotransmitter receptors improves the generative models of intracranial cortical dynamics. (A) Example spatial maps of gradients of four neuroreceptor densities and of the first four principal components interpolated from 44 regional averages. (B) Results of the principal component analysis across receptor densities and brain regions. (C) Parameter estimates for neuronal population parameters covary with the receptor density principal components 1-4 shown left as spatial projection. (D) Comparison between models that differ in which aspects of receptor density maps inform their inversion: single main excitatory and inhibitory receptor types, combinations of main excitatory and inhibitory receptor types, or combination of n principal components across all receptor types. Insert: histograms of free energy values obtained from single runs for each null model set (N1 – receptor correlation preserved, N2 – region boundary preserved, N3 – random) using randomised regressor values of the winning model (PC4); deltas indicate the difference between the winning model and the best null model in each set after 40 randomisations. Abbreviations: PC – principal component; Models: ANGa – PEB model with AMPA, NMDA, GABAA receptor densities as regressors; PC1-6 – PEB with 1 to 6 principal components as regressors; Baseline – standard CMC; N1-3 – Null models with differently shuffled (randomised) four principal components as regressors

Second, we assessed whether low dimensional representations across all 15 receptor densities improve fit compared to the excitatory / inhibitory systems models. Principal component analysis was applied to identify axes of maximal regional variance across the 15 receptor densities (Figure 2B, Supplementary Figure 1). Then, one to six of the resultant principal components were included in this second hierarchical PEB model. The winning model (*PC4*, including components one to four; Figure 2C) had a free energy difference of >5 compared to the previous winning model *ANGa* (posterior probability >.99) (Figure 2D). In fact, all models with more than one principal component outperformed the excitatory / inhibitory neuroreceptor models. This suggests that the principal components contained additional information — from the entire regional receptor density composition — which is relevant to explain neuronal dynamics, but which was not included in the densities of predominant E/I neurotransmitter systems.

Lastly, we evaluated whether the effects of receptor density information on model evidence could be explained away by spatial factors rather than by the distributions of receptor data. Since the receptor density estimates are spatially smooth and neighbouring cortical areas are likely to show more similar iEEG signals than those further apart, the addition of random but spatially smooth priors may improve model evidence. This was tested by inverting three null model sets through constrained shuffling of the receptor density values for each brain region: *N1*, shuffling while preserving correlation between regressors (receptor densities) within individual brain regions; *N2,* shuffling while preserving brain region boundaries; and *N3,* randomly shuffling of all receptor density values amongst iEEG channels individually. As expected, some of these null models outperformed an uninformed baseline model, but all had a significantly lower model evidence than the original winning models and any of the PCA models (Figure 2D).

### 2.3 Normative, receptor density informed neural mass models generalise

The previous steps demonstrated a statistical relationship between receptor density maps and oscillatory brain activity mediated through the parameters of a hierarchical model. Subsequently, we tested whether the informed neural mass model parameters generalise to other datatypes, tasks, and models; namely, do they have predictive validity? Specifically, the parameter posteriors of the winning model (*PC4*) were incorporated in a DCM of the well-studied mismatch negativity paradigm (59–61).

Mismatch negativity (MMN) is an auditory evoked potential that represents the difference of brain responses to repeated versus novel sounds. As a reference, we used a single participant EEG dataset that has been used previously to demonstrate dynamic causal modelling (59): model type and source locations (five-node distributed auditory network), recording modality (high density scalp EEG), and signal domain (time domain) differed from our normative iEEG dataset (Figure 3A, 3B). The iEEG normative synaptic parameter estimates from *PC4* served as regionally specific priors for intrinsic synaptic parameters of each cortical EEG source, i.e., neural mass model of the MMN. To identify an appropriate spatial mapping between the iEEG and EEG data modalities, we assigned prior values to the MMN network sources using averages of normative parameters which were estimated through Gaussian spatial kernels with varying widths (σ=1cm to 10mm) centred at their respective source locations (Figure 3A, 3C). In addition — to improve robustness of the model fits and to avoid local minima solutions — we iteratively fitted with multiple randomised initial conditions for each Gaussian kernel and selected the models with the largest free energies for Bayesian model comparison accordingly.

**Figure 3.**
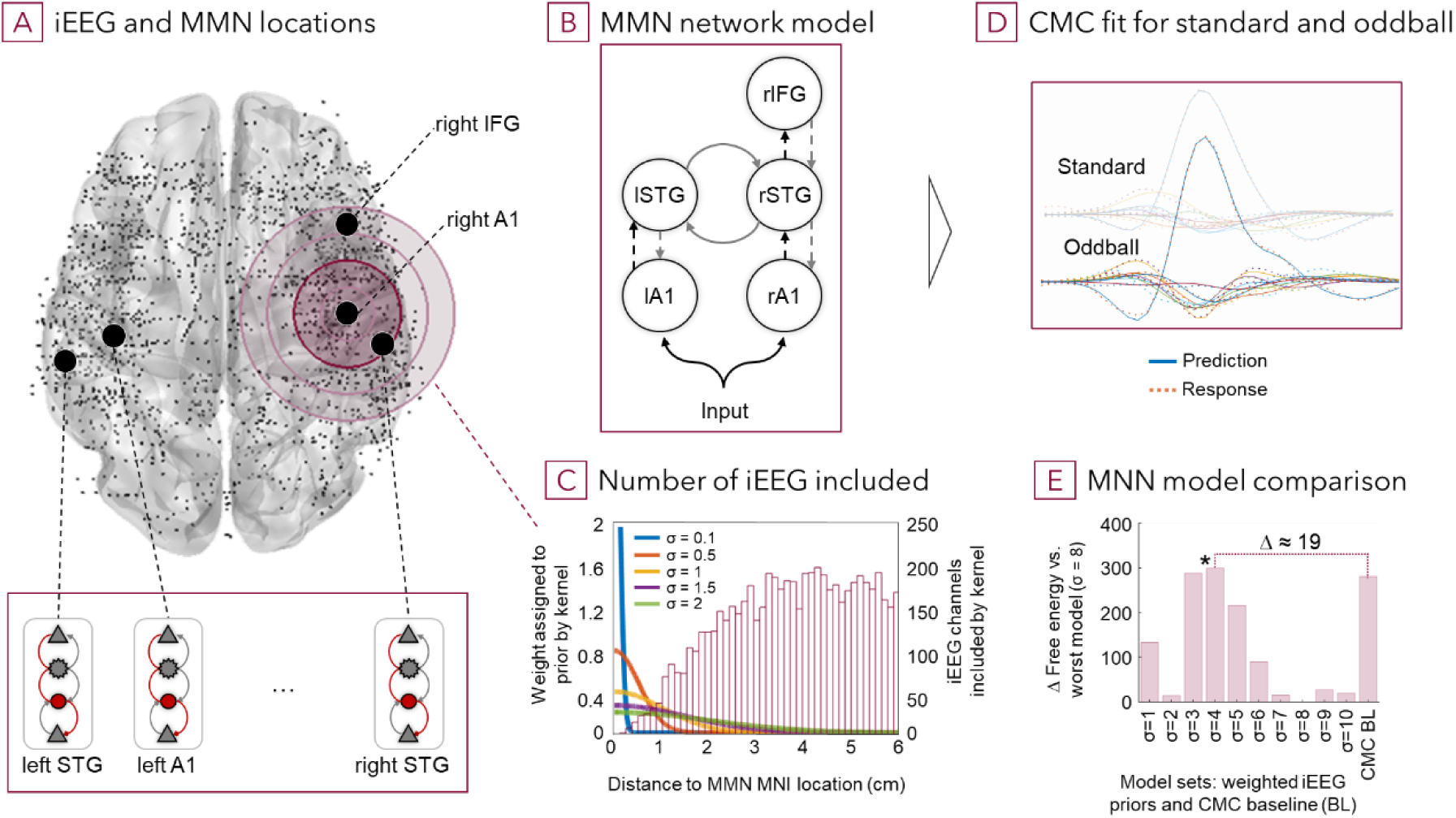
Receptor informed priors generalise to other modalities and signal domains. **(A)** Locations of the iEEG contacts (dots) for which normative parameters are available, and MMN regions of interest (black circles). The large red circles illustrate different spatial kernels. Insert: each MMN source is modelled with an individual CMC, **(B)** which are combined and included in the DCM network model with modulating forward and backward connections. **(C)** Euclidean distance was used to assign priors to the individual MMN source models using different Gaussian kernels to average the synaptic parameter priors from contributing (iEEG) locations. **(D-E)** An illustration of the ERP baseline and MMN response and prediction. Using different kernels and DCM parameter initiation values we compared the best models of each set: model evidence and fit were significantly improved versus the CMC model with standard priors.

Generalisability and predictive validity were confirmed, as receptor density informed normative (empirical) priors improved model evidence for the MMN dataset, despite differences in modality and domain of data features. Across the spatial kernels, there is a single peak in free energy at σ = 4cm (Figure 3E), indicating that this might be an appropriate scale of spatial averaging. The time-domain signals generated by the winning model had very good fit to the baseline and MMN response (Figure 3D) and the free energy difference compared to the uninformed model without receptor priors was >5 (posterior probability >.99).

## 3 Discussion

In this study, we demonstrated that neurotransmitter receptors densities shape resting state intracranial EEG dynamics. Normative receptor data and regional iEEG spectra were mapped to the parameters of a biophysically inspired generative model of cortical oscillations through an iterative Bayesian procedure. This resulted in an atlas of neurobiologically informed intracortical neuronal population synaptic parameters (i.e., intrinsic connectivity and rate constants), which generalise to DCM applications of independent datasets. These results speak to a central neuroscience question: how observable brain signals such as intracranial EEG power spectra can be explained by attributes of — and variations in — underlying neurobiology or structure (17, 62–64). Specifically, we provided quantitative evidence that neurotransmitter systems and their interactions underwrite emergent neuronal dynamics and that their influence can be captured with generative (neural mass) models. Additionally, we illustrated an efficient approach to translate across spatial scales in measures of brain structure and function in health and disorders.

### 3.1 Receptor densities shape regional electrophysiology

We focussed on assessing the explanatory power of cortical neurotransmitter receptors as key mediators of signal transduction between neurons (2, 65, 66). Aggregated regional receptor compositions represent a useful descriptor with molecular specificity, which may approximate meso- and macroscale electrophysiology (35–37). And topographic overlap of receptor density data and electrophysiological frequency bands was previously shown statistically (32). Therefore, the generative modelling approach presented here now offers mechanistic evidence that regional variability in receptor composition can explain differences in neuronal dynamics, when mediated through nonlinear models of population dynamics. It enabled us to find a parametric mapping of how spectral variation in low noise iEEG relates to the underlying regional composition of neurotransmitter receptor density and to obtain normative models of dynamics across the adult human brain.

### 3.2 Mesoscale models mediate between microscale synaptic properties and emergent macroscale brain dynamics

Regional receptor density compositions and iEEG signals are situated at the ‘mesoscale’: millimetre scale patches of cortical layers. The abstraction from individual microscopic structural components, such as neuronal cells, transmitters or receptors, allows to focus on emergent dynamics at population and circuit level which subsume underlying interactions such as modulatory or compensatory up- and downregulation of specific neuronal receptor subtypes (38). This means that quantitative representations of cortical dynamics such as power spectral densities can be aptly modelled using coupled neuronal populations: for instance, the DCM canonical microcircuit neural mass model. Furthermore, Bayesian inversion facilitates data-driven learning of the interrelation between receptor densities and CMC parameters. Employing this technique, we found that including entire receptor density compositions via principal components significantly improved models compared to incorporating only predominant excitatory and inhibitory neuroreceptors.

Further, changes in power spectral densities obtained from iEEG can be linked to CMC parameter variations, and PEB coefficients (Figure 2C) combined with PCA gradients (Figure 2A) allow to assess the influence of principal components onto regionally specific CMC parameters. But since principal components capture the shape of variation in normalised receptor densities and since the effects of the CMC parameter changes are accumulative, interpreting PSDs in terms of contributions of individual neurotransmitter receptors requires further testing. Overall, our results suggest that neuronal dynamics emerge from interacting neurotransmitter systems and are only partially determined by the excitatory-inhibitory balance (54). In essence, mesoscale models with limited but sufficiently complex parameter sets seemingly offer an appropriate trade-off between complexity and tractability for modelling regional and whole-brain dynamics.

### 3.3 Towards a normative atlas of synaptic population parameters of cortical dynamics

In addition to demonstrating mechanistic spatial dependencies between receptor densities and cortical dynamics, we provide a normative set of generative neural mass models across the cortical surface. Incorporating connectivity and biophysical properties of populations as well as additional regional receptor information, the ensuing synaptic parameter estimates represent a theoretically and practically relevant annotation of the human cortex, which may inform future dynamic causal modelling studies. Within the DCM framework, hidden neuronal sources — modelled as interacting populations — may be equipped with parameter priors and then fitted to EEG and MEG datasets in both time and frequency domains. Therefore, the parameter values that describe iEEG data here are useful to inform models of scalp EEG and MEG dynamics of different cortical populations. We illustrated this application by taking a previously validated DCM study of mismatch negativity as an example (59). Contextualised broadly, our approach — and the availability of the attending cortical parameter atlas — extends decade long modelling of brain dynamics with neural mass models (62, 63).

Incorporating the regional chemoreceptor composition into neural mass models to explain measured electrophysiology provides an opportunity to understand brain dynamics in terms of bottom-up and top-down causation. We envision that this allows to establish further links between chemoarchitecture and (clinically) measurable electrophysiology both in health and disorders. Consequentially, it may facilitate, for example, modelling and understanding of the entangled effects pharmacological compounds exert on different neurotransmitter systems. Our neurobiologically informed CMC seems suited to study these complex interactions and their electrophysiological signatures, as it accounts for the interrelation of 15 neuroreceptors, while generating larger-scale signals. Therefore, questions of how a drug affects (human) brains regionally or entirely through modulation of the underlying transmitter systems can be studied and explained. Hence, this approach complements biologically detailed computational models in drug discovery research as it abstracts from molecular and cellular mechanisms (68, 69) and instead models the consolidated effects compounds exercise onto the brain regionally or globally. In other words, it models characteristic emergent phenomena (70) taking various spatial scales and details into account.

### 3.4 Limitations and Constraints

The computationally efficient implementation within DCM allows to evaluate various hypotheses or explanations effectively. However, this comes at the expense of the Variational Laplacian model fitting approach being susceptible to locally — but not globally — optimal solutions (71, 72). In addition, since parameters collapse various underlying neuronal mechanisms, one-to-one reverse mapping requires elaborate testing procedures and might only provide approximate answers, potentially limiting predictive validity; for example, for explaining effects of pharmacological interventions without further data. This constitutes a general problem of inverse modelling at any scale and points to debates about scientific knowledge discovery (73), which are beyond this work. Other limitations of this study relate to the normative data used, as both the iEEG and the autoradiography datasets constitute only relatively small samples on which inference on normative electrophysiology and chemoarchitecture was drawn. However, future studies might further refine our findings and integrate additional prior information iteratively in similar model fitting procedures.

### 3.5 Empirically informed multimodal models of brain dysfunction may aid personalised medicine and novel therapeutic approaches

We have demonstrated an approach that uses Bayesian inference to link multimodal spatial datasets through generative models of brain function. This flexible methodology has wide-ranging applicability in neuroscience research; especially, since the capacity to evaluate brain (dys)function with complementary modalities at varying spatial scales increases (74–81). In the future, this might have practical implications as it could help to establish in-silico models to predict changes in whole-brain dynamics following therapies with molecular targets. Such models would aid in stratifying patients into aetiological groups according to electrophysiological phenotypes and support decision making in empirically informed, personalised interventions for brain dysfunction.

## 4 Methods

### 4.1 Overview

For this study, three sets of data and computational approaches were used (Figure 4). First, to model power spectral densities (PSD) for previously collected intracranial EEG (iEEG) data (50) we employed an established neuronal mass model, the canonical microcircuit (CMC) (76–80) (Figure 4A), and inferred its parameters using dynamic causal modelling (DCM) (52–55, 82). Subsequently, in a second level analysis with parametric empirical Bayes (PEB) (57, 58, 83) we tested which combinations of regional neurotransmitter receptor densities (RD) (54) (Figure 4B) improve model evidence (i.e., variational free energy), across all individually fitted regional CMC models. Finally, generalisability of the derived normative CMC synaptic parameter priors was assessed using independent mismatch negativity (MMN) data (59) and a case-adjusted modelling procedure to incorporate the informed priors (Figure 4C).

**Figure 4.**
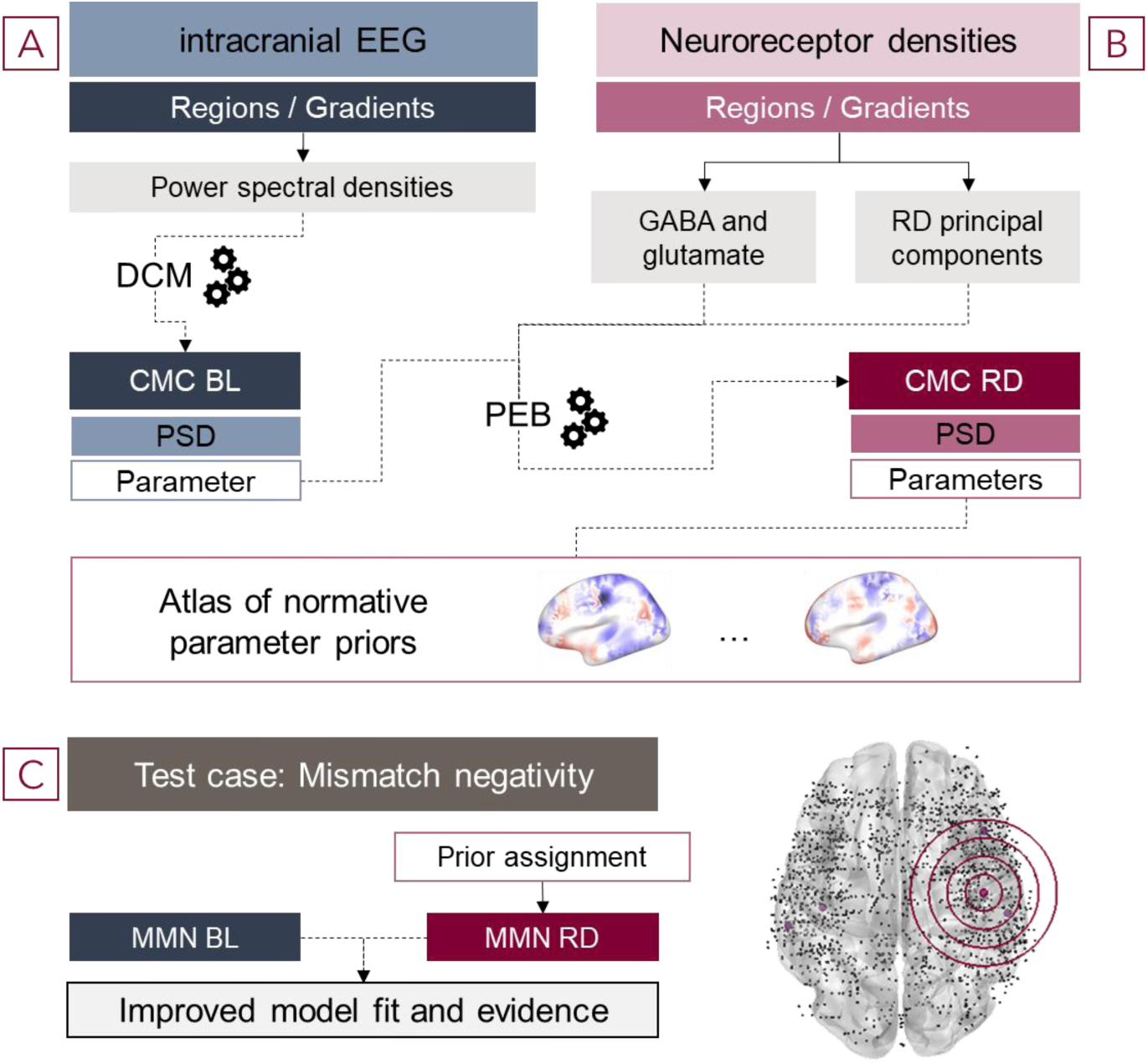
Modelling approach to integrate cortical dynamics and normative neuroreceptor density maps. **(A)** Interictal iEEG traces without overt pathological appearances are collated across 106 patients and 1770 contacts. Trace characteristics were summarised as power spectral densities (PSD) and individual four population models (canonical microcircuit model (CMC)) of cortical dynamics were fit to each individual contact’s trace using a variational Bayesian model fitting approach (dynamic causal modelling (DCM)). **(B)** Density profiles of 15 neurotransmitter receptors in 44 cortical were incorporated as covariates in a hierarchical analysis with parametric empirical Bayes (PEB) to assess if and how different combinations of receptors improve evidence. Two sets of hypotheses were tested: a) including only four major GABAergic and glutamatergic receptors, b) capturing regional variation of all receptors with principal component analysis and using these components. Three sets of randomised null models were used to validate results. We obtained neurobiologically (RD) informed CMC priors from this step. **(C)** To evaluate if the obtained priors generalise across modalities and signal domains, we used them in an unrelated test case (mismatch negativity (MMN)) and EEG traces in the time domain.

### 4.2 Normal-appearing intracranial EEG Data

The neurophysiological dataset used comprised 1770 artifact-free single-channel traces of iEEG time series (1512 from SEEG recordings, 258 from subdural grids/strips), recorded during 60 seconds of wakefulness with closed eyes (82, 83) (Figure 1A, 1B). These recordings were taken from 106 adult patients with drug-refractory epilepsy (54/52 male/female, mean age: 33.1+/-10.8 years) who underwent pre-surgical evaluation. Electrode contact locations were registered to a normative standard space ICBM152 (84, 85) with 38 regions, and on average, coverage was 0.9 channels/cm^2^ cortical surface, and 2.7 channels/cm^3^ cortical grey matter.

For this first normative atlas of iEEG (84), strict inclusion criteria relating to tissue type (MRI confirmed healthy tissue), location (peri-implantation imaging with CT/MRI), recording categorisation (additional control iEEG), conditions (standardized, resting wakefulness EEG with eyes closed), and sampling rate (minimum 200Hz) were applied. The iEEG recordings showed lobe-, and partially region-specific power spectral densities (PSD) with no statistically significant differences between power spectra of homologous regions in cerebral hemispheres.

Unmodified iEEG data as obtained from the website were used (86)

### 4.3 Neurotransmitter Receptor Density Autoradiography Data

Layer-specific density data of 15 neurotransmitter receptors (85, 86) was available for 44 regions (87, 88). The data were obtained using an in vitro receptor autoradiography protocol (89) on three neurologically healthy donor brains (2/1 male/female, 72-77 age range) (88). Region-specific balances of receptor densities (90) were estimated, showing, for example, that glutamate and GABA receptor densities vary substantially between cortical regions (45, 89).

Since RD correlations between layers (supra-, infra-, granular) were on average r=.83 (80% percentile range: [.62, .99]) across receptors, we used normalised unweighted averages of the three layers as regressors in our PEB analyses.

### 4.4 Mapping iEEG and Receptor Density Data

We mapped unihemispheric receptor density sample locations onto the MNI space of the iEEG data. Subsequently, the iEEG contact locations were projected onto the left hemisphere – into the same space as the receptor density data – as no statistically significant hemispheric differences were previously reported for either dataset (45, 91).

### 4.5 Dynamic Causal Modelling

Dynamic causal modelling (52–55, 82) is a Bayesian computational framework that facilitates computationally efficient Variational Laplacian fitting of generative models to empirical data. Iterative inversion with variational Bayes under Laplace assumption (92) fits model parameters by gradient descent along a variational free energy bound on log model evidence. In this formulation, free energy balances accuracy and complexity, penalising models with multiple redundant free parameters that do not significantly improve model fit. The fitted models contain means, variances and covariances of model parameters (multivariate normal probability density over parameter posteriors). DCM is available as part of the open source statistical parametric mapping (SPM12) academic software, in which several neural mass model types, and inference and analyses methods are implemented (93).

The CMC (92) is an extension of the Jansen and Rit model (94). Four populations, superficial pyramidal, spiny stellate, inhibitory interneuron and deep pyramidal, with population/layer-specific excitatory, inhibitory, and modulatory coupling strengths and time constants, constitute a canonical cortical microcircuit (Figure 1E). The CMC parameters and their priors are biophysically informed (50, 95), and population dynamics are described by differential equations which include a linear, pre- to post-synaptic potential, and a sigmoid, post-synaptic to action potential, operator. Typically, these biophysical population models describe electrical activity (membrane currents and potentials) on the cortical surface, and a separate, observation model is applied, e.g., a leadfield model that projects the activity of sources to scalp EEG sensors or local field potentials in the case of iEEG. Steady state iEEG activity is modelled by fitting parameters to power spectral (or cross-spectral) density summaries of oscillations. Effectively, this uses a local linear perturbation analysis around the fixed point of a nonlinear model (55).

For each of the 1770 iEEG recordings, an individual CMC was fitted to the observed power spectral density, which delivered full posterior parameter estimates in the 14 neuronal population parameters (three excitatory and three inhibitory between population coupling strengths, four inhibitory/self-modulatory strengths, and four population time constants), and a free energy estimate for each CMC. These fully fitted models were the basis for subsequent hierarchical modelling steps. Note that this assumes the iEEG data can be regarded as generated by a single canonical (i.e., normative) subject.

### 4.6 Parametric Empirical Bayes

Parametric Empirical Bayes (PEB) is a hierarchical Bayesian modelling approach (57, 58, 83) used for second (group or between DCM) level analysis and parameter estimation based on first level DCM CMC parameter posteriors. Here, PEB was employed to test hypotheses via changes in model evidence (free energy at the second level) (i) after including neuroreceptor densities as explanatory variables (i.e., GLM regressors) in the PEB model, and (ii) about relations between neuroreceptor densities and first level CMC parameters. Additionally, hierarchical models tend to provide more robust parameter estimates than first level parameters as they are constraint by group effects and are less likely to get stuck in local minima during model inversion.

To evaluate the effect of RD data on model evidence two PEB approaches were employed. The first PEB included densities of four candidate receptors (AMPA, NMDA, GABA_A_ and GABA_B_) as regressors to analyse the effect of each receptor type on model evidence (free energy). This resulted in a 1770 x 4 matrix (1770 iEEG channels, four normalised receptor densities) constituting the model space and enabled us to evaluate 15 different models (combinations of covariates): four models with a single receptor density, six with two regressors, four with three regressors and one model with all four receptor densities. We hypothesised that a mix of major excitatory and inhibitory receptor densities would increase model evidence significantly.

For the second PEB, we first performed a principal component analysis (PCA) to capture cortical topographic variation and clusters of normalised receptor densities in reduced dimension. This was helped by receptor fingerprint similarity of neighbouring, connected or functionally comparable areas (37) and by spatially cooccurring receptor expression (Figure 2B, Supplementary Figure 1). Different features of the underlying receptor data are reflected in principal component gradients, for example, the distinctively low density of most receptors in the motor cortex is apparent in PC1 and higher expression of receptors such as GABAa, NMDA, M2, alpha2, D1, 5-HT2 in early visual areas in the occipital lobe is discernible in PC2 (Figure 2A). The first one to six principal components (PC) of the RD-region matrix, which together explain 91% of variability (PC1-6 explained .405, .245, .106, .073, .048, .034 respectively) of the 15 receptor densities across 44 regions, were used as regressors. This resulted in a 1770 x 6 matrix with PC covariates, and for this second PEB we combined the covariates in order from the first to the sixth principal component. Thus, we inverted six second level models, used Bayesian model comparison and the softmax operator to obtain posterior probabilities over models. We reasoned that the entire population dynamics can only partially be explained by the excitatory/inhibitory receptors alone and that variability in the composition of all regional receptor densities is informative.

Furthermore, to assess if improvements in model evidence are due to non-specific spatial coherence among parameters, we used three independent sets of null models which included the first four principal components to validate results of the overall winning model (*PC4*). For the first null model set (*N1* – receptor correlation preserved), values of all receptors together were randomly exchanged between regions which preserved the correlation of densities within a region. In the second null model variant (*N2* – regional boundary preserved), receptor values were exchanged between regions for each regressor individually, thus, it was only ensured that iEEG channels within a region have the same values for a given regressor. The last null models (*N3* – random) were unconstrained within a regressor, therefore, values of a receptor density were randomly shuffled between iEEG channels, which implies that channels in the same region likely have different receptor values. With the first null model set, the association between regional densities and observed dynamics was tested. With the second, the contribution of regional composition, i.e., correlations between densities, was evaluated. And with the third, additionally, the importance of within-region consistency was assessed. For each of the null model variants, regressor values were shuffled 40 times, delivering a representative number of individual model inversions per set for comparison.

### 4.7 Mismatch Negativity

The mismatch negativity (MMN) is an event-related potential (ERP) component evoked by detectable violations in acoustic regularity (60). It was extensively investigated with DCM (59, 61, 96, 97) and therefore, it served as our test study for generalisability beyond the dataset from which our normative parameter values were estimated (98). The reference models and data were obtained from Garrido et al. (2007) (59). The dataset comprised a 128-channel single subject EEG with inter-stimulus (-100 to 400ms) 512Hz recordings, down-sampled to 200Hz and band-pass filtered between 0.5 and 40Hz.

In accordance with the original model, we included five sources: left and right primary auditory cortex (A1) and superior temporal gyrus (STG), and right inferior frontal gyrus (IFG) (Figure 3A); and assumed that forward and backward connections between hemispheric sources as well as between within-source populations can be modulated (Figure 3B). Instead of the three population DCM ERP neural mass model in the original paper (96), we applied the four population DCM CMC with Hanning-filter for fitting the EEG data in the time-domain. To inform these CMC models, the normative excitatory and inhibitory coupling, and modulatory parameter priors were used. Normal distribution probability density functions (PDF) with different standard deviations (σ=1 to 10) functioned as spatial kernels to assign weighted iEEG RD-grounded posteriors as DCM intrinsic coupling parameter priors to the five MMN source locations (Figure 3A, 3C). In addition, different starting values (expectations) for the expectation-maximisation algorithm (variational Bayes) were set to avoid local extrema solutions. For this, random initiation values were sampled from the range of coupling parameter values across the five MMN sources. Thus, for each of the ten spatially weighted coupling prior sets eighteen CMC-RD using different random coupling parameter initiation values were inverted. For comparison, eighteen baseline CMC (CMC-BL) were also inverted using standard DCM CMC priors as well as parameter initiation values of the winning CMC-RD to ensure comparability. The CMC-RD models and the CMC-BL model with the highest evidence within their set were subsequently selected for comparison and visualisation (Figure 3E).

## 5 Data, Materials, and Software Availability

Functions and models can be found in the statistical software package for neural imaging and electrophysiological data ‘Statistical Parametric Mapping’, Version 12 (93) implemented in MathWorks MATLAB.

---

**Supplementary Figure 1.**
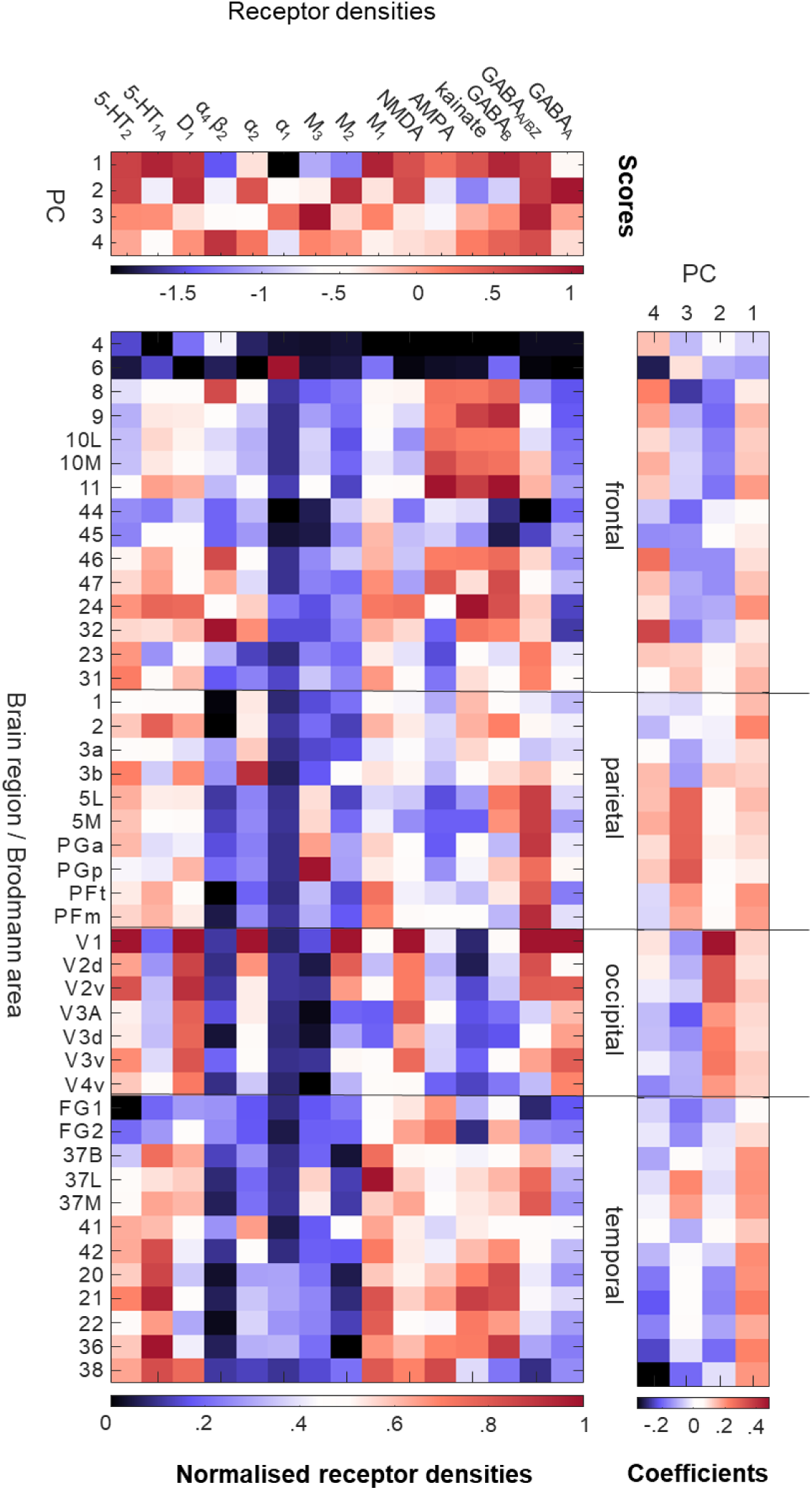
Principal component analysis. Additional detail for the PCA across receptor densities and brain regions (see also Figure 2B) with labels for brain regions, lobes, and receptor densities. Region labels were named in accordance with (37).

